# The automated eukaryotic pangenome pipeline EukPan reveals accessory genome differentiation beyond core-gene phylogeny in *Aspergillus oryzae*

**DOI:** 10.64898/2026.09.13.751290

**Authors:** Kiyohiko Seki, Masatoshi Goto, Taiki Futagami, Yukio Nagano

## Abstract

Pangenome analysis reveals recurrent gene-content variation beyond a single reference genome, but its application to eukaryotes is constrained by inconsistent gene annotation. ANNEVO predicts gene models from genome FASTA assemblies without RNA-seq data. We developed EukPan, an automated post-annotation pipeline that standardizes GFF/GTF files, selects representative isoforms, constructs proteomes, infers orthogroups, builds a concatenated single-copy core-protein alignment, and summarizes shared accessory orthogroups while excluding orthogroups detected in only one genome. Applied with ANNEVO to 123 *Aspergillus oryzae* genomes, EukPan identified 11,245 core and 4,407 shared accessory orthogroups. The core-protein phylogeny broadly recovered the reported A-H classification, whereas accessory-genome analyses clearly separated the 33 group-A strains from the other 90 strains. Directional analysis identified 62 group-A-associated and 158 group-A-depleted orthogroups, with major facilitator superfamily (MFS) transporter and fungal Zn_2_Cys_6_ transcription-factor domains prominent in the depleted set. Among 93 orthogroups present in all non-A strains and absent from all group-A strains, 59 mapped to 10 segments of RIB40, the standard *A. oryzae* reference genome and a non-A (group-F) strain. EukPan therefore enables reproducible, coordinated core– and accessory-pangenome analysis from eukaryotic genome assemblies.

## Introduction

Pangenome analysis provides a framework for characterizing the full gene repertoire of a species or a closely related group of organisms. Since the microbial pangenome concept was formalized, gene repertoires have commonly been divided into core genes shared by most or all genomes and accessory genes present in only a subset of genomes [1]. Core genes are useful for reconstructing conserved evolutionary relationships, whereas accessory genes reflect gene gain, loss, duplication, and lineage-specific retention. Accessory-gene content can therefore reveal ecological, metabolic, and phenotypic diversity that is not represented by a single reference genome [2].

For bacteria, a widely used sequence-to-pangenome approach is the Prokka–Roary workflow. Prokka annotates assembled genomes and produces GFF3 files suitable for direct input into Roary, which clusters annotated coding sequences into core and accessory gene sets [3–4]. Bakta and DFAST provide more recent alternatives for standardized bacterial genome annotation [5–6], whereas Panaroo and PPanGGOLiN offer alternative downstream pangenome frameworks incorporating graph-based error correction or explicit partitioning of gene families [7–8]. We recently applied the Prokka–Roary workflow to *Bacillus subtilis* and found that accessory-gene profiles grouped an Indian bekang isolate with the Japanese natto clade, despite this relationship being less evident in the core-genome phylogeny [9].

Establishing an equivalent routine workflow for eukaryotes has been more difficult because eukaryotic protein-coding genes contain introns, may produce multiple transcript isoforms, and require structural gene prediction before protein comparison. Evidence-driven pipelines such as BRAKER3 can generate high-quality gene models when RNA-seq and protein evidence are available [10], but matched transcriptomic data are unavailable for many assemblies available through the International Nucleotide Sequence Database Collaboration (INSDC), which links the DNA Data Bank of Japan (DDBJ), the European Nucleotide Archive (ENA), and GenBank at the National Center for Biotechnology Information (NCBI). Recent deep-learning and ab initio approaches, including Helixer and ANNEVO, can predict eukaryotic gene models directly from genome sequences without requiring RNA-seq data [11–12], making uniform annotation of large genome collections increasingly feasible.

Gene prediction alone, however, does not yield an analysis-ready pangenome. Bacterial workflows such as Prokka–Roary cannot simply be transferred because Roary was designed for prokaryotic coding sequences. Eukaryotic annotations are supplied in heterogeneous GFF/GTF formats and may include multiple isoforms, requiring standardized conversion, representative-isoform selection, protein extraction, orthogroup inference, construction of single-copy core alignments, and summarization of accessory-orthogroup distributions. A reproducible post-annotation workflow connecting these steps is therefore needed for large-scale eukaryotic pangenome analysis.

The conserved and variable components of a pangenome provide complementary views of diversification. Core-gene phylogenies describe relationships inferred from conserved genes, whereas accessory-gene distributions capture recurrent gene-content variation arising through gain and loss, duplication, introgression, mobile-element activity, and lineage-specific retention. In fungi, variable gene repertoires can include functions involved in secondary metabolism, transport, secreted-enzyme activity, and environmental responses [2], potentially revealing strain differentiation that is not apparent from conserved-gene phylogenies.

*Aspergillus oryzae*, the yellow koji mold widely used in food fermentation, provides a suitable system in which to compare these complementary genomic perspectives. The RIB40 reference genome established a foundation for *A. oryzae* genomics [13]. Watarai et al. defined clades A-H as those containing Japanese industrial *A. oryzae* strains used for products including sake (Japanese rice wine), miso (fermented soybean paste), shoyu (soy sauce), and mirin (sweet rice seasoning); industrial use did not fully determine the clades [14]. Their large-scale comparison also identified domestication-associated evolution and mosaic genome structure. A comparative pangenome study of *A. oryzae* and *Aspergillus flavus* further characterized broad patterns of gene-content similarity and metabolic repertoires across the two species [15]. However, it remained unclear whether the A-H classification inferred from core genes would correspond to accessory-genome structure in a larger collection of *A. oryzae* strains.

To address these computational and biological questions, we developed EukPan, an automated post-annotation pipeline for eukaryotic pangenome analysis. In the workflow implemented here, assembled genomes were quality-screened using BUSCO [16] and annotated with the RNA-seq-free tool ANNEVO. EukPan then standardized the resulting GFF/GTF gene models, selected representative isoforms, generated proteome datasets, inferred orthogroups, and produced both core-protein alignments and accessory-orthogroup presence-absence matrices. Application of the combined ANNEVO-EukPan workflow to 123 *A. oryzae* genomes broadly recovered the established A-H classification in the core-gene phylogeny, while identifying group A as the dominant axis of accessory-genome differentiation. Directional functional summaries, evidence-filtered homology review, and locus-level comparisons between the group-F reference strain RIB40 and the group-A strain TK-34 were then used to characterize recurrent group-level differences.

## Results

### EukPan implements 12 post-annotation steps for core– and accessory-genome analyses

Figure 1 summarizes the automated workflow used in this study. Following BUSCO-based quality screening [16] and RNA-seq-free gene prediction with ANNEVO [12], EukPan accepts corresponding genome FASTA and GFF/GTF annotation files and implements 12 post-annotation steps. It (1) standardizes and converts GFF, GFF3, or GTF annotations using AGAT [17], (2) retains the longest isoform for each gene, (3) extracts protein sequences using gffread [18], (4) simplifies protein FASTA headers using SeqKit [19], and (5) runs OrthoFinder [20].

**Figure 1.**
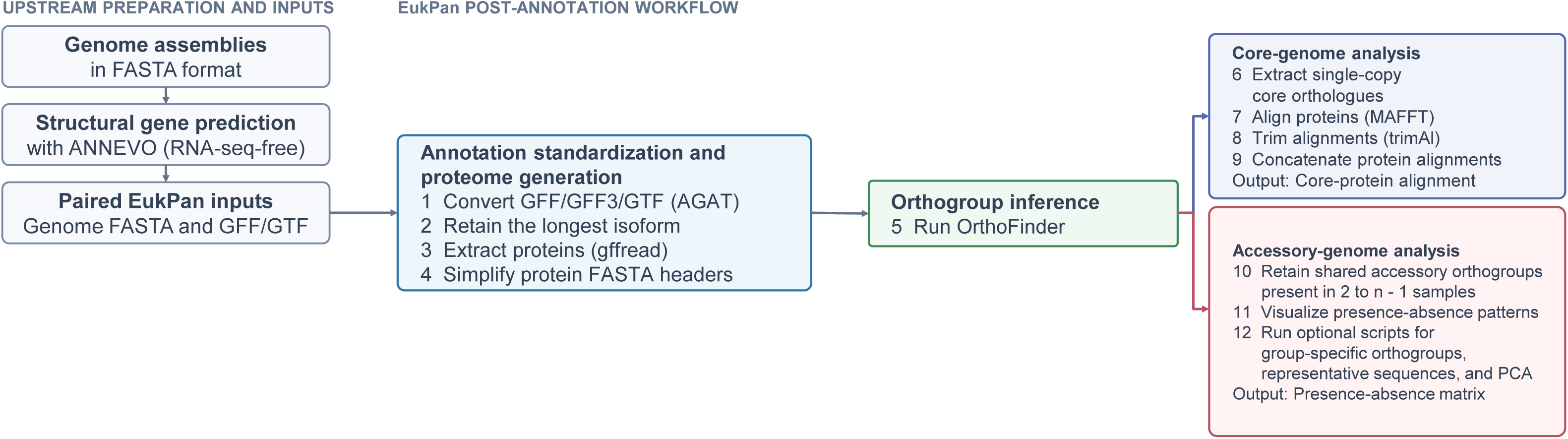
Overview of the ANNEVO-EukPan workflow. ANNEVO predicts structural gene models directly from genome assemblies in FASTA format without matched RNA-seq data. The resulting GFF/GTF annotation files are paired with the corresponding genome FASTA files and supplied to EukPan. Steps 1-11 form the core workflow: (1) conversion and standardization of GFF, GFF3, or GTF annotations using AGAT, (2) retention of the longest isoform for each gene, (3) protein-sequence extraction using gffread, (4) simplification of protein FASTA headers, (5) OrthoFinder analysis, (6) extraction of single-copy core orthologues, (7) sequence alignment using MAFFT, (8) alignment trimming using trimAl, (9) concatenation of trimmed single-copy core-protein alignments, (10) extraction of shared accessory orthogroups, and (11) visualization of accessory-orthogroup presence-absence patterns. Step 12 comprises optional scripts for group-specific orthogroup analysis, representative-sequence extraction, and PCA. Shared accessory orthogroups were defined as non-core orthogroups detected in at least two but fewer than all genomes.

From the OrthoFinder output, EukPan (6) extracts single-copy core orthologues, (7) aligns them using MAFFT [21], (8) trims the alignments using trimAl [22], and (9) concatenates the trimmed single-copy core-protein alignments. In parallel, it (10) extracts shared accessory orthogroups and (11) visualizes accessory-orthogroup presence-absence patterns. These first 11 steps form the core EukPan workflow. Step 12 comprises three optional downstream analyses implemented as helper scripts: group-specific orthogroup analysis, representative-sequence extraction, and PCA (principal component analysis) of accessory-orthogroup presence-absence profiles.

### Application of EukPan to 123 *Aspergillus oryzae* genomes

We applied EukPan to 123 *Aspergillus oryzae* genomes. All 126 BUSCO-screened assemblies, including the 123 retained for EukPan analysis and the three excluded assemblies, are listed with genome accession numbers, complete BUSCO scores, A-H group assignments where applicable, and post-screening inclusion status in Supplementary Table S1. The dataset comprised 33, 5, 18, 2, 16, 13, 9, and 3 strains assigned to groups A-H, respectively, together with 24 strains without an A-H assignment. Orthogroups detected in only one genome were excluded from the shared-accessory matrix. EukPan identified 11,245 core orthogroups present in all 123 strains and retained 4,407 shared accessory orthogroups present in at least two but fewer than all strains, corresponding to a prevalence range of 2-122 strains. The validated machine-readable presence-absence matrix and summary files are provided in Supplementary Data S1. All subsequent accessory-genome visualizations, distance calculations, clustering analyses, and PCA were based on these 4,407 shared accessory orthogroups. The number of shared accessory orthogroups present in each genome ranged from 1,662 to 2,053, with a mean of 1,864.

### The core-gene phylogeny broadly recovered the previously reported groups

ModelTest-NG selected VT+I+G4+F for the amino acid substitution model [23]. Using this fixed model, we reconstructed a maximum-likelihood phylogeny in IQ-TREE 3 from the concatenated single-copy core-protein alignment [24]. The resulting phylogeny broadly recovered the previously reported A-H classification of *A. oryzae* strains [14] (Fig. 2).

**Figure 2.**
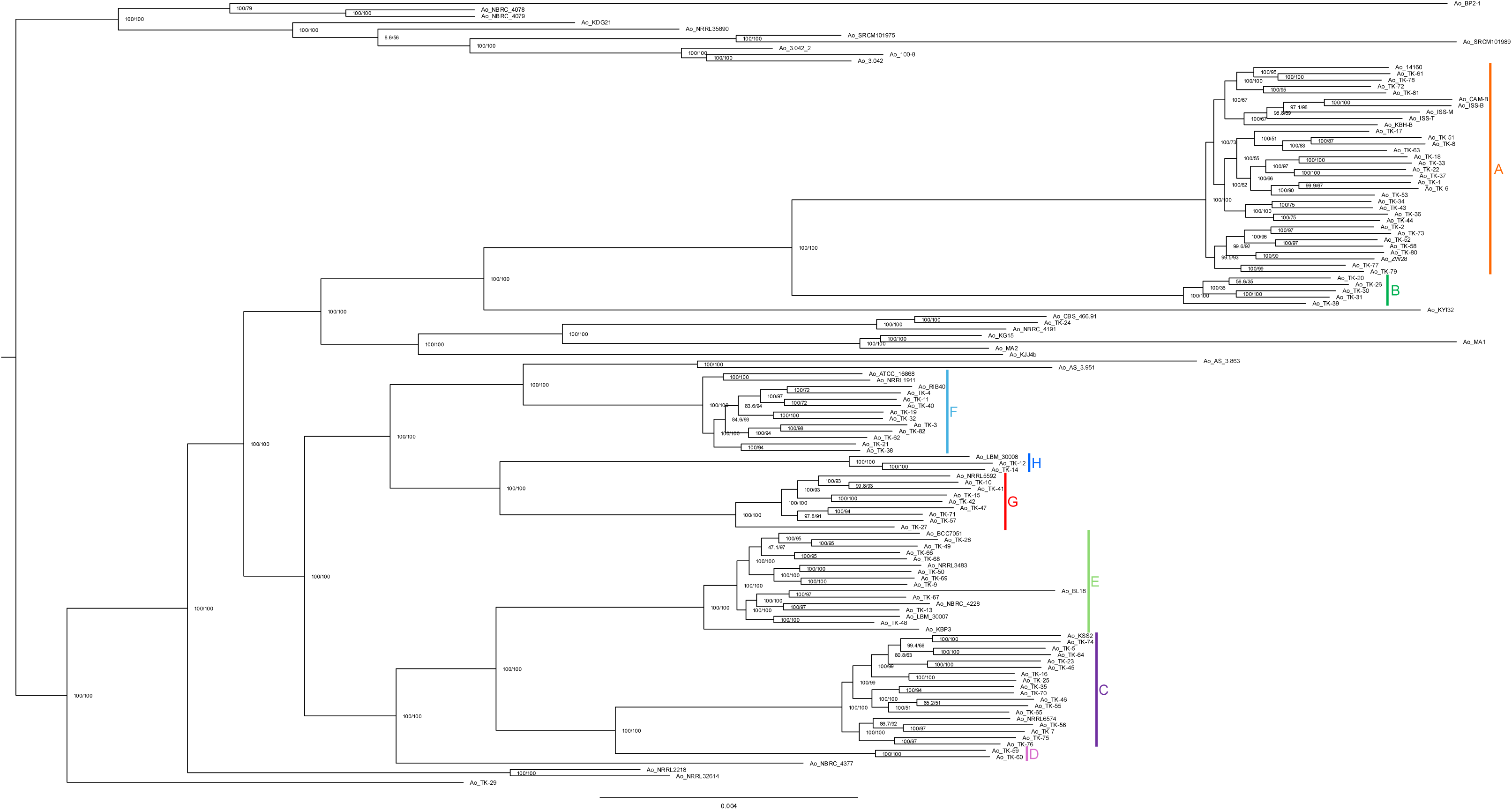
Maximum-likelihood phylogeny based on single-copy core proteins. The amino acid substitution model was evaluated using ModelTest-NG, and the tree was inferred using IQ-TREE 3 from a concatenated alignment generated from 10,535 trimmed single-copy core-orthologue alignments from 123 *Aspergillus oryzae* strains. The fixed VT+I+G4+F model was used. The resulting phylogeny was midpoint-rooted and visualized using FigTree v1.4.4. Node labels show SH-aLRT support and ultrafast bootstrap support percentages, respectively, based on 1,000 replicates each. Colored brackets indicate the previously defined groups A-H. Strains without a previous group assignment are not included within a colored bracket.

### The core-gene split network contained a central box-like structure

To further examine phylogenetic signals in the core-gene alignment, we constructed a NeighborNet network in SplitsTree App using uncorrected p-distances [25] (Supplementary Fig. S1). The resulting network comprised 156 splits, was classified as cyclic, and had a reported fit value of 98.4%. It contained a central box-like structure rather than displaying an entirely tree-like arrangement.

### Shared accessory-orthogroup prevalence was strongly nonuniform

The prevalence of the 4,407 shared accessory orthogroups varied widely across the 123 strains, ranging from 2 to 122 strains per orthogroup (Fig. 3). Of these orthogroups, 1,366 were present in 10 or fewer strains, whereas 608 were present in 120 or more strains. The largest individual peaks occurred at prevalences of two strains, comprising 422 orthogroups, and 122 strains, comprising 394 orthogroups. An additional peak at 90 strains comprised 101 orthogroups; its relationship to the group-A/non-A separation is examined below.

**Figure 3.**
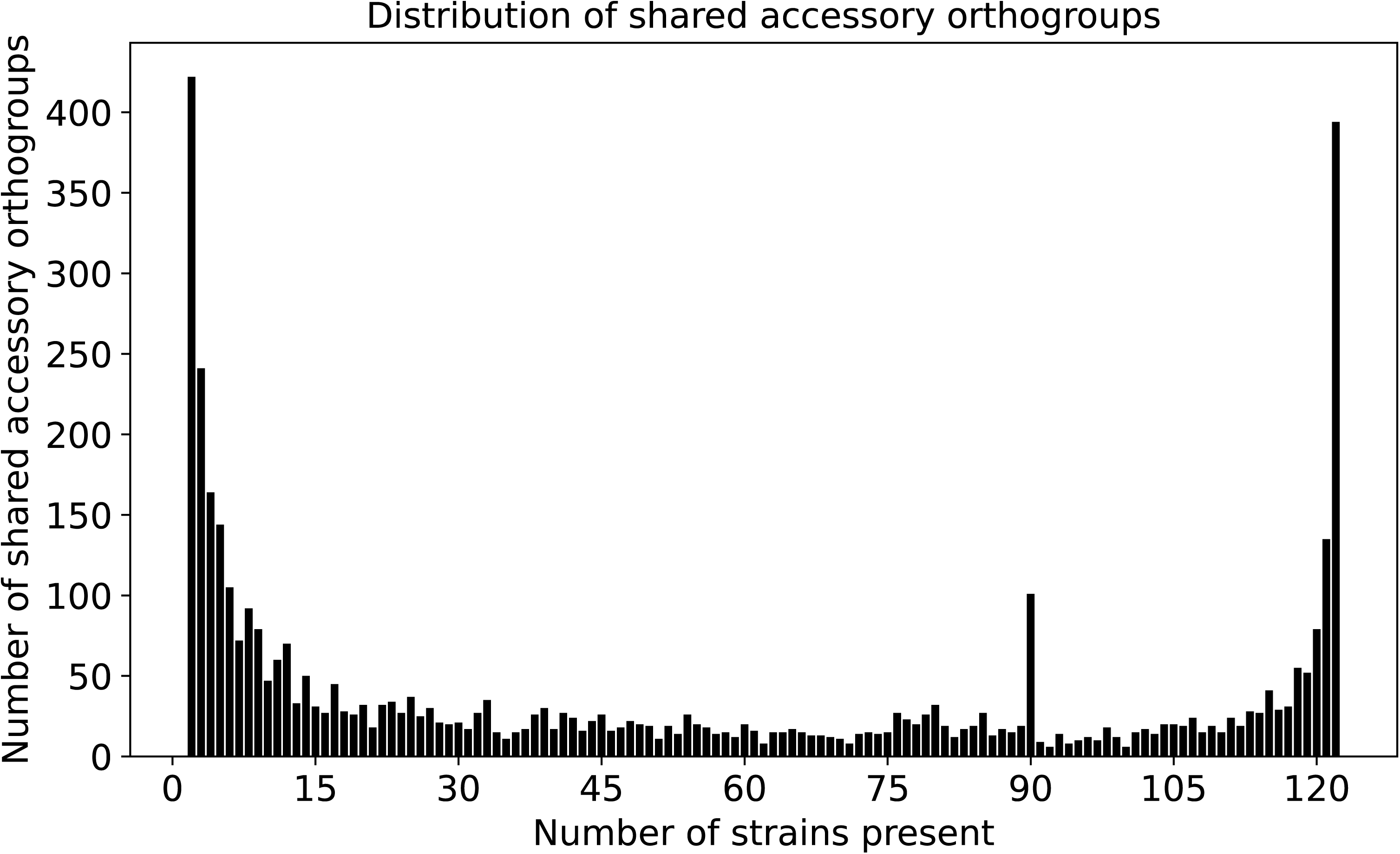
Distribution of shared accessory orthogroups among the 123 strains. The horizontal axis shows the number of strains in which each orthogroup was present, and the vertical axis shows the number of orthogroups at each prevalence value. The analysis included 4,407 non-core orthogroups detected in 2–122 strains. Prominent peaks occurred at prevalence values of 2, 90, and 122 strains.

### Accessory-genome PCA clearly separated group A from the other strains

Principal component analysis of accessory-orthogroup presence-absence profiles clearly separated group A from all remaining strains along the first principal component (PC1) (Fig. 4). PC1 explained 23.66% of the total variance, whereas PC2 explained 7.00%. PC1 scores ranged from 15.903 to 17.492 among group-A strains and from –9.281 to 1.189 among the remaining strains. The nearest scores from the two sets were therefore separated by 14.714 units. PC2 captured additional variation among the remaining strains but did not disrupt their separation from group A along PC1.

**Figure 4.**
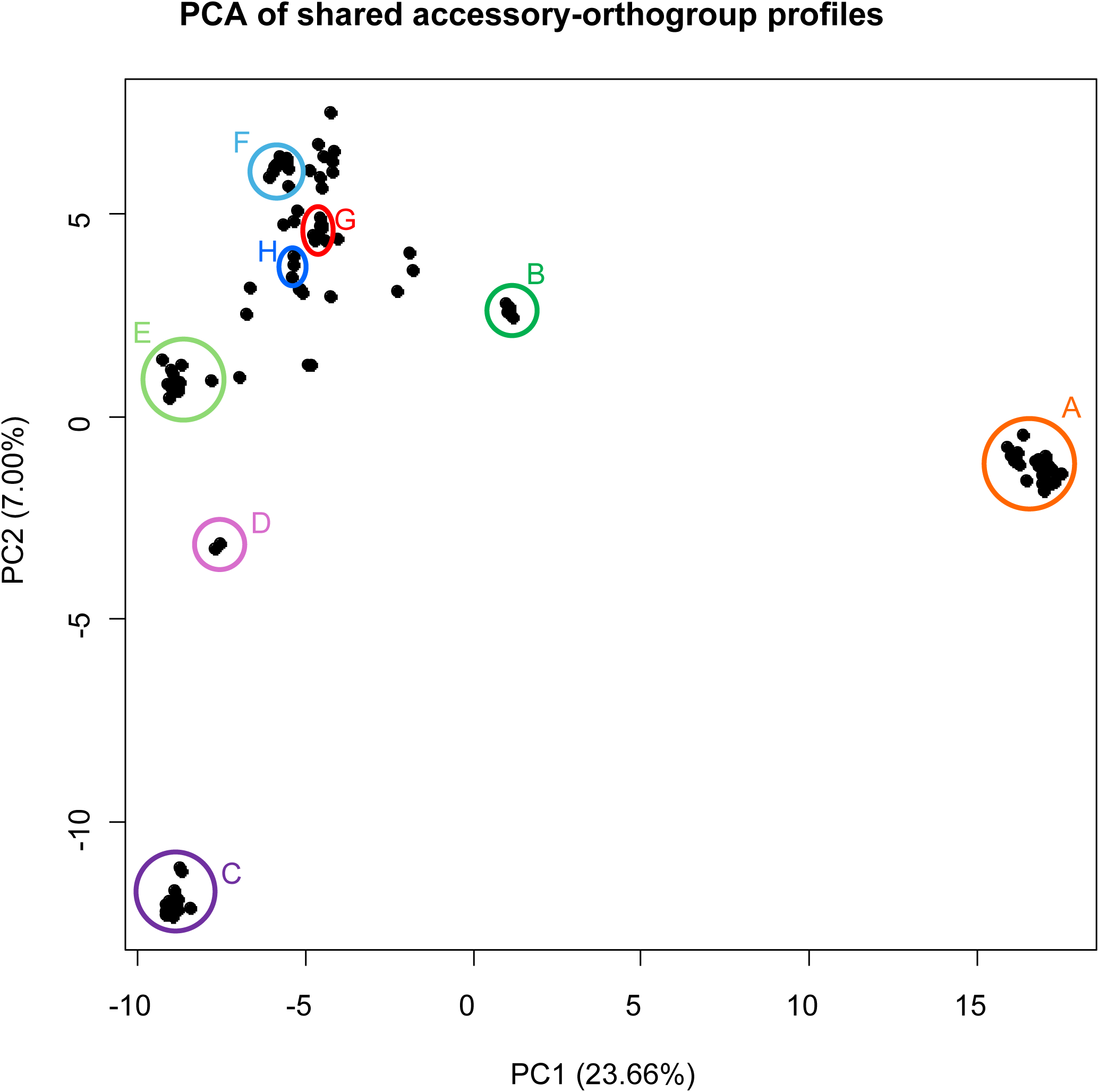
Principal component analysis of shared accessory-orthogroup presence–absence profiles. Each black point represents one strain; colored outlines and labels identify the previously defined groups. PC1 and PC2 explained 23.66% and 7.00% of the total variance, respectively. Group labels were assigned after PCA for visualization and were not used to calculate the principal components.

### Distance-based analyses confirmed group-A accessory-genome differentiation

Jaccard-distance-based analyses of accessory-orthogroup presence-absence profiles showed the same overall pattern. In the hierarchically reordered distance matrix, all 33 group-A strains formed a contiguous block (Fig. 5). The mean pairwise Jaccard distance was 0.217 within group A, 0.481 between group A and the remaining strains, and 0.355 within the remaining set.

**Figure 5.**
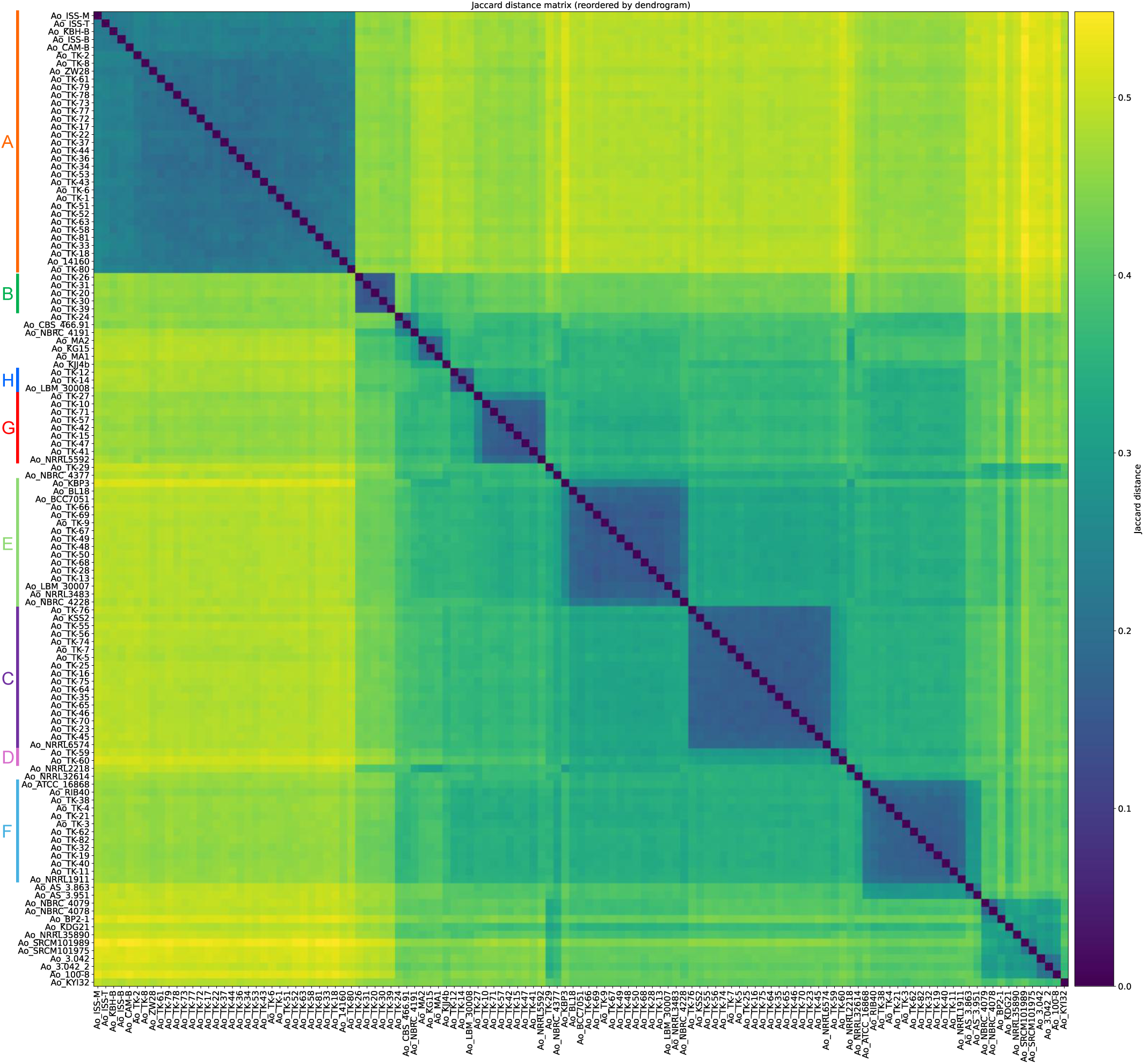
Jaccard-distance matrix based on shared accessory-genome profiles. Rows and columns represent the same 123 strains and were reordered according to average-linkage hierarchical clustering with optimal leaf ordering. Lower Jaccard distances indicate greater similarity in shared accessory-orthogroup content. Colored brackets indicate the previously defined groups A-H.

The presence–absence heatmap and hierarchical clustering similarly recovered group A as a coherent accessory-genome cluster (Fig. 6). The remaining strains showed additional substructure that corresponded only partly to the groups recovered from the core-gene phylogeny. For visual comparison, the core-gene tree was also displayed with the root placed on the branch leading to group A (Supplementary Fig. S2). This rooting was used solely for visualization and does not imply that group A represents a biological outgroup. The displayed core-gene tree topology did not fully correspond to the accessory-genome dendrogram.

**Figure 6.**
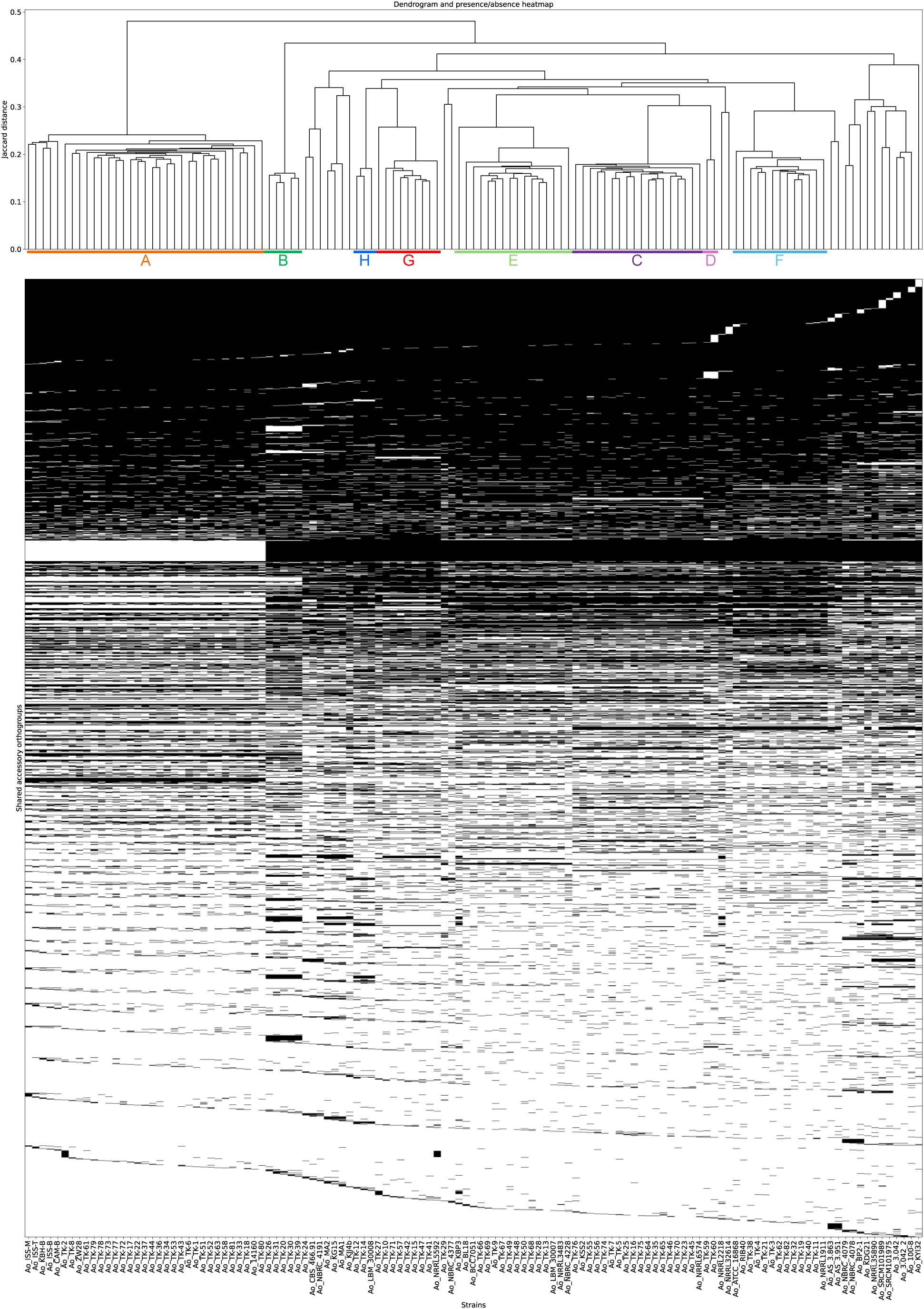
Hierarchical clustering and presence–absence heatmap of the shared accessory genome. The upper dendrogram was constructed from Jaccard distances using average-linkage hierarchical clustering with optimal leaf ordering. Columns represent the 123 strains arranged in dendrogram order. Rows represent the 4,407 shared accessory orthogroups, ordered first by prevalence and then by their binary presence–absence patterns. Black indicates presence, and white indicates absence. Colored bars indicate the previously defined groups A-H.

Of the 101 orthogroups at the 90-strain peak in Fig. 3, 93 were present in all 90 non-A strains and absent from all 33 group-A strains. We refer to this all-versus-none subset as strict group-A-depleted orthogroups, where strict denotes the prevalence pattern rather than a separate orthology class (Supplementary Table S2, ‘Strict A-depleted’ worksheet). Thus, the peak was largely attributable to the group-A/non-A separation.

### Directional group comparisons identified an asymmetric functional signature of group A

To identify recurrent accessory orthogroups associated with the previously defined strain groups, we used the optional EukPan group-comparison scripts to define two directional sets for each of groups A-H. Group-associated orthogroups were present in at least 90% of the focal group and at most 10% of the complementary non-group set, whereas group-depleted orthogroups met the reciprocal thresholds. The non-group complement included all other strains, including the 24 strains without an A-H assignment. The numbers of group-associated and group-depleted orthogroups, respectively, were 62 and 158 for group A, 100 and 57 for B, 31 and 17 for C, 28 and 40 for D, 34 and 12 for E, 11 and 12 for F, 19 and 14 for G, and 84 and 22 for H (Supplementary Table S2 and Supplementary Fig. S3). Because group sizes ranged from 2 to 33 strains, these counts are most stable for the larger groups and should be interpreted cautiously for groups B, D, and H.

Group A showed the largest absolute difference between the two directional counts, with 96 more group-depleted than group-associated orthogroups (158 versus 62). The 158 group-A-depleted orthogroups included these 93 strict orthogroups, which generated most of the prevalence peak at 90 strains in Fig. 3. Combining OmicsBox and InterPro-derived Gene Ontology (GO) annotations yielded at least one GO term for 44 of 62 group-A-associated orthogroups (71.0%) and 111 of 158 group-A-depleted orthogroups (70.3%); InterPro domains were assigned to 49 of 62 (79.0%) and 132 of 158 (83.5%), respectively. After OmicsBox and InterPro-derived GO assignments were combined, Biological Process (BP), Cellular Component (CC), and Molecular Function (MF) terms were represented in 27, 16, and 43 of the 62 group-A-associated orthogroups, respectively, and in 48, 17, and 97 of the 158 group-A-depleted orthogroups, respectively. The two sets therefore had similar overall annotation coverage, although this does not exclude differences in gene-prediction accuracy or functional-annotation bias. GO and InterPro annotations were summarized as orthogroup frequencies to characterize the functional composition of the two directional lists (Fig. 7b,c).

**Figure 7.**
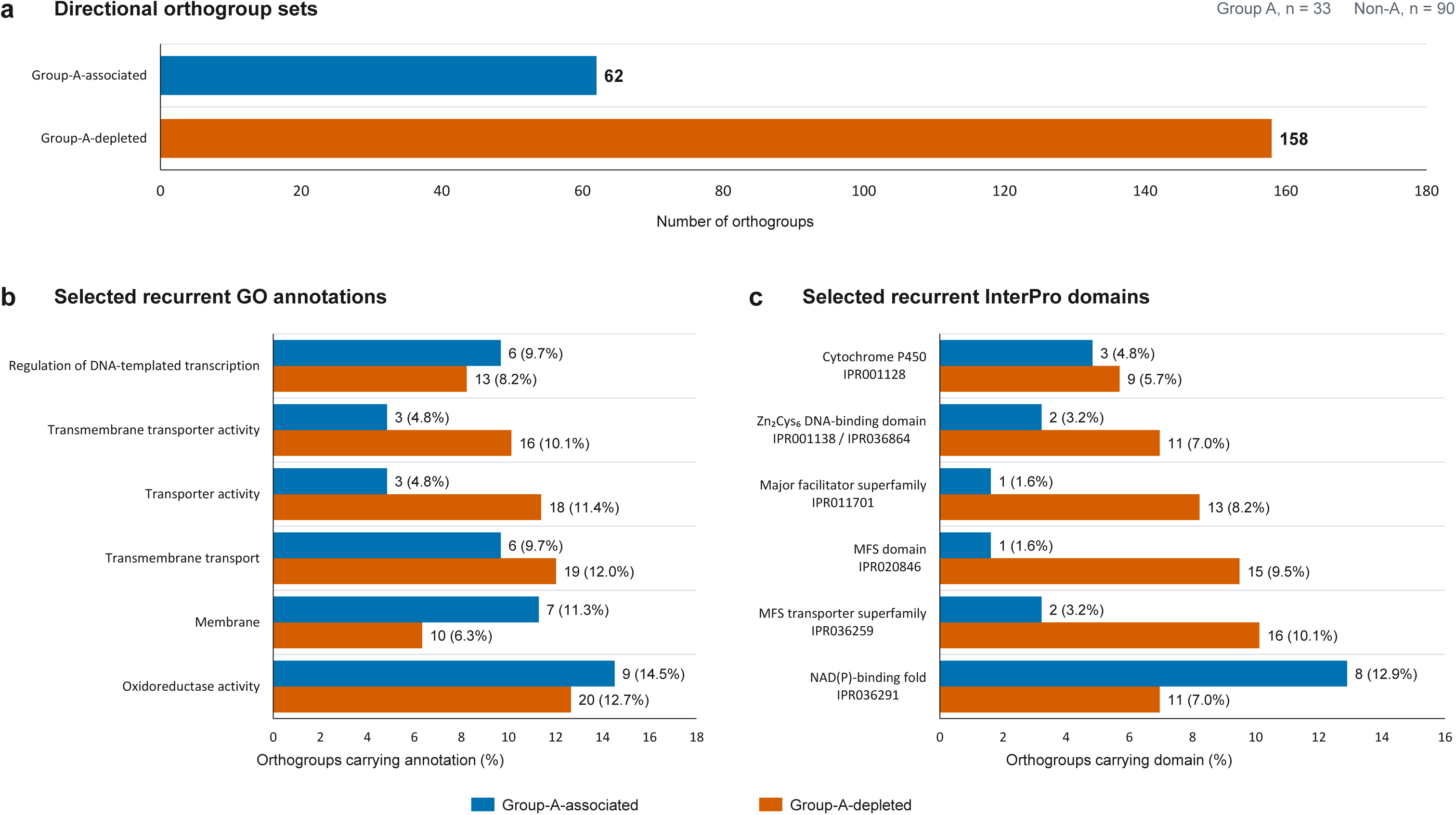
Directional functional profile of group. **A.** (a) Numbers of group-A-associated orthogroups, defined as present in 90-100% of group-A strains and 0-10% of non-A strains, and group-A-depleted orthogroups, defined using the reciprocal thresholds. (b) Frequencies of selected recurrent GO annotations in the two directional sets. (c) Frequencies of selected recurrent InterPro domains. Displayed annotations were selected to summarize prominent categories and contrasts between the lists. Percentages were calculated using all orthogroups in each directional set as the denominator (62 group-A-associated and 158 group-A-depleted orthogroups); labels show the orthogroup count and percentage. An orthogroup can carry multiple GO terms or InterPro domains. The panels characterize the annotated composition of the lists; no statistical hypothesis testing was performed.

Among group-A-associated orthogroups, the most frequent GO annotations were oxidoreductase activity (MF; 9/62), membrane (CC; 7/62), transmembrane transport (BP; 6/62), and regulation of DNA-templated transcription (BP; 6/62). In the associated set, the NAD(P)-binding fold was the most frequent InterPro annotation (8/62, 12.9%), followed by P-loop NTPase (4/62, 6.5%) and protein kinase-like and cytochrome P450 domains (3/62, 4.8% each) (Supplementary Table S2). Representative annotations included protein kinases, a Zn_2_Cys_6_ transcription-factor protein, cytochrome P450 monooxygenases, a tryptophan dimethylallyltransferase, nonribosomal peptide synthetase modules, and a MATE transporter. Group-A-depleted orthogroups frequently carried oxidoreductase activity (MF; 20/158), transmembrane transport (BP; 19/158), transporter activity (MF; 18/158), transmembrane transporter activity (MF; 16/158), and transcription-regulatory annotations (BP; 13/158) (Fig. 7b). Across all group-A-depleted orthogroups, 16 of 158 (10.1%) carried MFS transporter-superfamily annotations (IPR036259), 15 of 158 (9.5%) carried MFS-domain annotations (IPR020846), 13 of 158 (8.2%) carried major-facilitator-superfamily annotations (IPR011701), and 11 of 158 (7.0%) contained fungal Zn_2_Cys_6_ transcription-factor DNA-binding domains (IPR001138/IPR036864). Representative annotations included multiple MFS permeases, Zn_2_Cys_6_ transcription-factor proteins, P450 proteins, and nonribosomal peptide synthetase modules (Supplementary Table S2). The selected cross-set comparison in Fig. 7c shows that the NAD(P)-binding fold was proportionally more frequent in the associated set, whereas MFS and Zn_2_Cys_6_ DNA-binding domains were proportionally more frequent in the depleted set.

### Evidence-filtered sequence analysis identified eight group-A-depleted homologs of experimentally characterized proteins

Representative proteins from the two group-A directional sets were compared with reviewed *Aspergillus* proteins and the leading matches were evaluated using targeted literature searches. To limit functional overinterpretation, only candidates classified in the high-evidence category were linked to the functions of experimentally characterized homologs. Eight group-A-depleted orthogroups met this criterion, whereas no group-A-associated orthogroup did.

Six of the eight high-evidence group-A-depleted orthogroups were absent from all 33 group-A strains and present in all 90 non-A strains. Their best reviewed matches were AflM/Ver-1, AflN, AflL, AvfA, OmtB, and OmtA, with 89.0-99.6% amino acid identity over 88-100% of the query length. Experimental studies support the functions of corresponding homologs [26–31]. For the AflL-like candidate OG0012552, the experimental anchor was *Aspergillus nidulans* StcL [28], a separate match with 80.8% identity and 98% query coverage, rather than a direct functional test of its best hit, *Aspergillus parasiticus* AflL. The best reviewed matches and literature-supported homologs are distinguished in Supplementary Data S1. Because high sequence identity alone does not establish biosynthetic activity, these assignments indicate high-confidence homology rather than production of a particular metabolite.

Two additional group-A-depleted orthogroups had high-evidence matches. OG0012523 was absent from all group-A strains and present in all non-A strains and matched *A. oryzae* CYP505C3 at 97.3% identity and 99% query coverage; CYP505C3 is a self-sufficient cytochrome P450 [32] whose fatty acid and alcohol hydroxylation activities have been characterized biochemically [33]. OG0012739 was absent from group A and present in 82 of 90 non-A strains and matched the *A. oryzae* heptelidic-acid-cluster regulator HepR at 100% identity and coverage [34].

### Strict group-A-depleted orthogroups were concentrated in multiple RIB40 regions

To examine their genomic organization, we used the group-F strain RIB40 (GCA_000184455.3) as the reference and the group-A strain TK-34 (GCA_009686785.1) as the representative comparison genome; both group assignments and accessions are listed in Supplementary Table S1. All 93 strict group-A-depleted orthogroups were present in all 90 non-A strains and absent from all 33 group-A strains. All 93 orthogroups mapped to RIB40, and 83 occurred in 15 compact genomic clusters. Comparison of each cluster plus 20-kb flanks with the TK-34 assembly identified 10 clusters that met the operational criteria for a two-flank deletion in TK-34. These 10 regions contained 59 of the 93 orthogroups and represented 189,888 bp of RIB40 sequence (Fig. 8a). Orthogroup-level coordinates and TK-34 comparison classifications are provided in Supplementary Table S2 and Supplementary Data S1.

**Figure 8.**
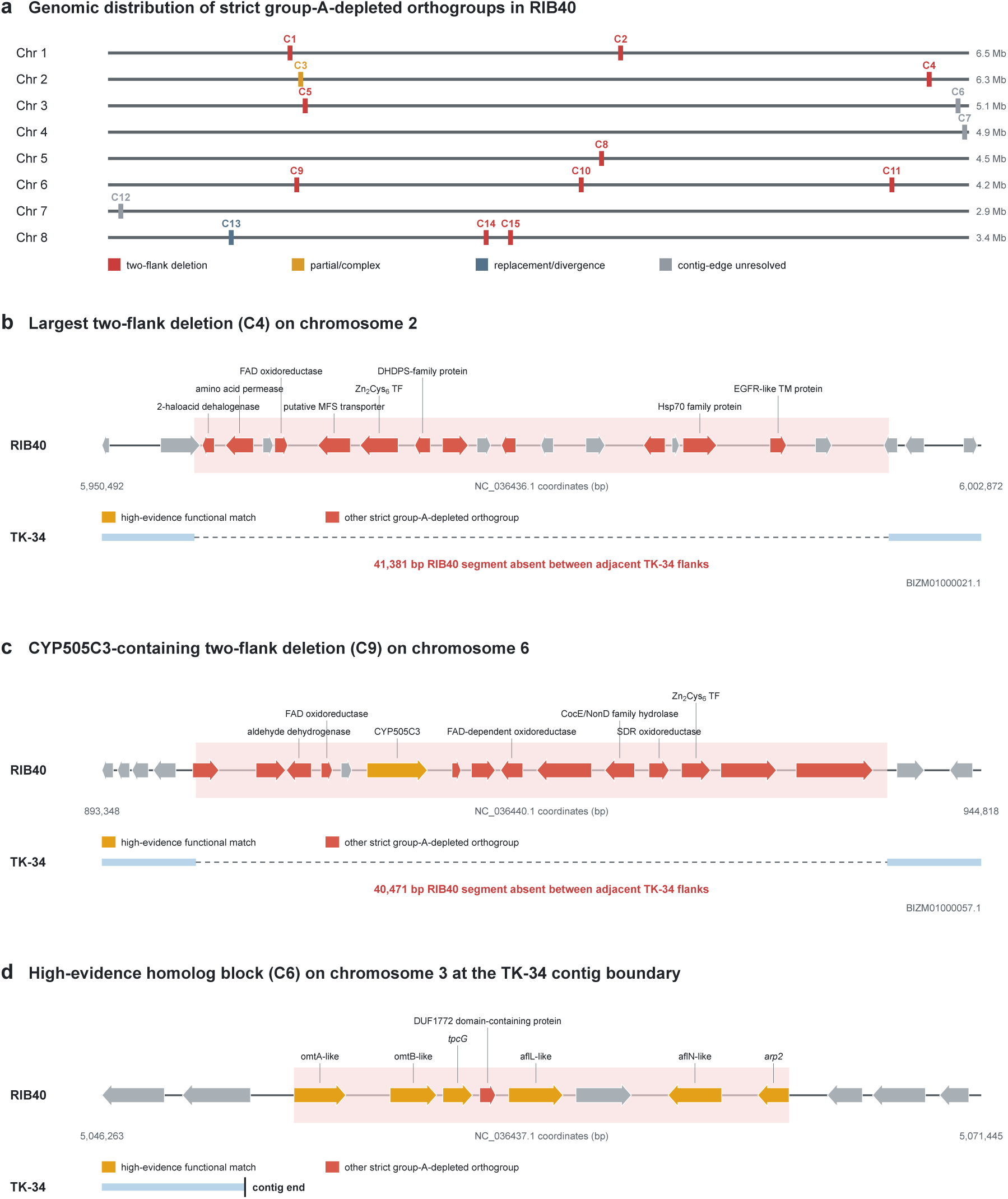
Genomic organization of strict group-A-depleted orthogroups in the group-F strain RIB40 and comparison with the group-A strain TK-34. (a) Locations of 15 compact RIB40 clusters containing 83 of the 93 orthogroups present in all 90 non-A strains and absent from all 33 group-A strains. Chr 1-Chr 8 denote RIB40 chromosomes 1-8, with chromosome lengths shown at right. Red marks indicate 10 two-flank deletions in TK-34; orange, blue-gray, and gray marks indicate partial/complex, replacement/divergence, and contig-edge-unresolved regions, respectively. The 10 two-flank deletions contained 59 of the 93 strict group-A-depleted orthogroups and totaled 189,888 bp in RIB40. The contig-edge-unresolved clusters C6, C7, and C12 lie entirely within 100 kb of the right end of chromosome 3, the right end of chromosome 4, and the left end of chromosome 7, respectively. (b) The 41,381-bp C4 interval on chromosome 2. (c) The 40,471-bp C9 interval on chromosome 6, which contains CYP505C3. (d) The 14,183-bp C6 high-evidence homolog block lies 57,739-71,921 bp from the right end of RIB40 chromosome 3 and was classified as subtelomeric; the TK-34 alignment terminates at a contig end, so a physical deletion cannot be bracketed by two flanks. Gene models and labels follow the RIB40 RefSeq GenBank annotation GCF_000184455.2. Established common names were used when supported; otherwise RefSeq gene, product, or locus-tag names were retained. Gold arrows indicate high-evidence functional matches, red arrows indicate other strict group-A-depleted orthogroups, and gray arrows indicate neighboring genes. The orthogroup absence pattern was defined across all group-A strains, whereas nucleotide-level comparison in this figure used TK-34 as one representative group-A genome.

The two largest two-flank deletions were a 41,381-bp region on chromosome 2 containing 14 strict group-A-depleted orthogroups (C4; Fig. 8b) and a 40,471-bp region on chromosome 6 containing 16 such orthogroups, including CYP505C3 (C9; Fig. 8c). Three clusters lay entirely within the 100-kb subtelomeric interval used in a comparative analysis of *Aspergillus* section *Flavi* [35]: C6 on chromosome 3, 57,739-71,921 bp from the right end; C7 on chromosome 4, 20,602-27,733 bp from the right end; and C12 at positions 25,245-61,362 near the left end of chromosome 7 (Fig. 8a). Six of the eight high-evidence homologs were colocated within C6 in a 14,183-bp region (Fig. 8d). The corresponding RIB40 RefSeq annotations include the established names *tpcG* and *arp2* for two genes. For C6, C7, and C12, conserved alignments could not be identified on both sides because the corresponding TK-34 alignments reached contig boundaries. Of the remaining clusters, one showed a partial or complex deletion and one was consistent with sequence replacement or strong divergence rather than a simple deletion.

### Group E contained the largest set of strains with documented use in soy sauce production

Industrial-use information provided additional context for the group-wise comparison. Among groups A-H, strains with documented use in soy sauce production occurred in groups D, E, and H, whereas the available metadata did not identify a consistent specific use for the other groups [14]. Groups D and H each contributed only two strains with documented use in soy sauce production, making group-level interpretation uncertain. By contrast, group E comprised 16 strains, including 10 with documented use in soy sauce production. In the available metadata, no use other than soy sauce production was reported for group E, and use information was not available for the remaining six strains. Group E therefore provided the most informative set for examining accessory-gene characteristics potentially shared among soy-sauce-associated strains, while recognizing that documented industrial use did not define every strain in the group.

EukPan identified 34 group-E-associated and 12 group-E-depleted orthogroups (Supplementary Table S2). Each of the 34 associated orthogroups was detected in all 10 group-E strains with documented use in soy sauce production and in five or six of the six strains without use information. GO terms were assigned to 13 of the 34 associated orthogroups (38.2%), and InterPro domains to 22 (64.7%). Descriptive annotations included several candidates compatible with secondary-metabolism or transport functions, including nonribosomal peptide synthetase-related and ABC-transporter features. BLASTP identified matches to reviewed *Aspergillus* proteins for 13 associated and four depleted orthogroups, but none met the high-evidence criterion used to assign an experimentally characterized function. These group E results therefore define a reproducible group-level accessory signature without assigning any individual orthogroup a validated soy-sauce-related function.

Among the 12 group-E-depleted orthogroups, seven were absent from all 16 group-E strains and five were detected in one group-E strain. Because the reviewed-protein matches for this set were incomplete or weak and none reached the high-evidence category, no specific pathway loss was inferred from these orthogroup distributions.

### Functional characterization of the other A-H groups

Other groups yielded more limited but potentially informative signatures. Group-C-associated orthogroups included hydrolase activity (8/31), RNA-interference/gene-silencing proteins, and a Dicer-like helicase, whereas group-G-associated orthogroups included GH18 and class V chitinases and glycosyltransferase-family proteins, suggesting recurrent differences in carbohydrate– and cell-wall-related functions.

Groups B and H had relatively many group-associated orthogroups but contained only five and three strains, respectively, and their annotations spanned oxidation-reduction, transport, transcriptional regulation, and secondary metabolism. Group F had only 11 associated orthogroups and low functional-annotation coverage. These patterns provide functional descriptions of the directional lists and nominate candidates for subsequent biological evaluation. Annotation coverage across all 16 directional sets is summarized in Supplementary Fig. S4.

## Discussion

EukPan bridges structural gene prediction and reproducible eukaryotic pangenome analysis. It standardizes annotations, selects representative isoforms, extracts proteomes, infers orthogroups, constructs single-copy core alignments, and summarizes recurrent accessory variation in one workflow (Fig. 1). Coupling EukPan with RNA-seq-free ANNEVO makes the same analysis practical for public genome assemblies that lack matched transcriptomes.

In *Aspergillus oryzae*, the core and accessory analyses provided complementary views of strain structure. The core-protein phylogeny broadly agreed with the A-H classification [14] (Fig. 2), but group A was much more sharply separated by accessory-genome composition in PCA, Jaccard-distance, and heatmap analyses (Figs. 4-6). The prevalence peak at 90 strains reflected 93 strict group-A-depleted orthogroups, 59 of which mapped to 10 RIB40 regions (Figs. 3 and 8). These results identify a spatially clustered component of the group-A-associated gene-content pattern; the contribution of these regions to the overall separation was not quantified.

Transport and transcriptional regulation were prominent in the group-A directional profiles (Fig. 7). MFS proteins mediate diverse fungal uptake and export processes [36], while Zn_2_Cys_6_ transcription factors regulate metabolic and stress-response programs [37] and often occur in secondary-metabolite gene clusters [38]. Six high-identity Afl homologs occurred in RIB40 cluster C6 and were absent from the group-A orthogroup profiles (Fig. 8d). This pattern is consistent with earlier evidence that loss or inactivation of aflatoxin-pathway genes contributes to aflatoxin nonproduction in *A. oryzae*, a property central to its use in food fermentation [39–42]. The TK-34 comparison ended at a contig boundary, however, so the underlying sequence change remains unresolved; retention of homologs in non-A strains does not imply aflatoxin production. CYP505C3, by contrast, lay in the fully bracketed C9 deletion (Fig. 8c), whereas HepR did not satisfy the strict all-versus-none pattern. These matches nominate experimentally anchored candidates but do not demonstrate pathway activity or fermentation phenotype.

Three RIB40 clusters, C6, C7, and C12, lie within 100 kb of chromosome ends (Fig. 8a,d). Species-specific genes and secondary-metabolite clusters are enriched near chromosome ends in *Aspergillus* section *Flavi*, where reduced synteny and rearrangement indicate elevated structural plasticity [35]. Their positions therefore provide a plausible context for recurrent gene-content turnover, but the present sample does not establish statistical enrichment. Because TK-34 alignments for these regions reached contig boundaries, chromosome-level group-A assemblies will be required to distinguish deletion from divergence, pseudogenization, or incomplete assembly.

Industrial koji performance depends strongly on the degradation of starch and protein substrates [14,43]. Three strict group-A-associated candidates were annotated as an amino acid transporter or aspartic-peptidase-related proteins (Supplementary Table S2), offering testable links to protein utilization. None met the high-evidence criterion, however, so their presence cannot be taken as evidence of greater proteolytic capacity. Brewing performance may also reflect allelic, copy-number, expression, and secretion differences in core genes.

Group E warrants separate consideration because 10 of its 16 strains had documented use in soy sauce production, more than in the other soy-sauce-associated groups [14]. Previous genomic and transcriptomic comparisons of soy sauce koji strains have emphasized proteolytic enzymes [44–45]. All 34 group-E-associated orthogroups occurred in those 10 strains, but also in most group-E strains of unknown use. Their secondary-metabolism and transport annotations did not meet the high-evidence criterion. Although 4-ethylguaiacol contributes to soy sauce aroma, its reported formation involves halotolerant yeasts [46], and our data do not link this reaction to group-E *A. oryzae*. These orthogroups characterize group E but cannot yet be assigned a role in soy sauce production; expression, metabolite, and gene-disruption studies are needed.

Other group-wise comparisons provided narrower hypotheses: group C included hydrolase and RNA-silencing candidates, while group G included carbohydrate-active enzymes. Results for groups B, D, and H are especially sensitive to their small sample sizes, and the group-F-associated set had low annotation coverage. These profiles should therefore be treated as candidate repertoires rather than evidence of specialized functions.

Excluding orthogroups detected in only one genome focused the analysis on recurrent variation and was intended to reduce sensitivity to isolated assembly and annotation errors. In eukaryotes, apparently genome-specific genes can result from fragmented assemblies, missed exons, split or fused models, alternative-transcript annotation, or isoform selection. The two-genome threshold consequently also excludes true single-strain genes and cannot remove errors shared across assemblies; the 4,407 shared accessory orthogroups are not an exhaustive catalogue of unique genes.

Previous fungal pangenome studies have linked accessory repertoires to lineages, habitats, infection, recombination, genome plasticity, and pathogenesis [2,47–50]. Hamanaka et al. recently identified Starships, giant transposable elements carrying cargo genes, as contributors to *A. oryzae* diversification during domestication. Cargo-gene differences relative to *A. flavus* and among *A. oryzae* strains implicated these elements in fermentation-related variation [51]. This provides a mechanistic context for accessory-genome diversity, although we did not test whether the regions identified here are Starship-associated. The expanded within-species *A. oryzae* analysis extends this work by showing that accessory content can reveal differentiation that is muted in a core-protein phylogeny. The NeighborNet network contained conflicting phylogenetic signals consistent with the mosaic history proposed for *A. oryzae* (Supplementary Fig. S1) [14], although a split network alone does not establish recombination. Its incomplete correspondence with the accessory dendrogram (Fig. 6 and Supplementary Fig. S2) underscores the value of examining both.

Several limitations remain. Results depend on assembly and structural-annotation quality; BUSCO screening cannot recover missing sequence or correct systematic gene-prediction errors [16]. Orthogroup inference may also be affected by paralogy, pseudogenization, fragmented models, and assembly gaps, while representative-protein annotations may not capture variation among all members. The RIB40-TK-34 comparison directly tested only one group-A assembly, and three regions ended at TK-34 contig boundaries (Fig. 8). GO and InterPro frequencies describe list composition rather than enrichment. The broader taxonomic search used for the group-H-associated set limits direct comparisons of annotation coverage, and even high-evidence homology does not establish activity. The multiple accessory-genome views summarize the same matrix and are not independent biological validations. This single-species demonstration establishes workflow feasibility, but comparative accuracy, runtime, and performance across eukaryotic lineages were not benchmarked.

Together, ANNEVO and EukPan converted quality-screened genome assemblies into coordinated core– and accessory-pangenome resources and revealed group-A differentiation that was substantially clearer in accessory content than in the core-protein phylogeny. The workflow supports reproducible reuse of public eukaryotic genomes while preserving explicit thresholds and machine-readable outputs.

## Methods

### Genome dataset, ANNEVO annotation, and EukPan execution

The proof-of-concept dataset comprised assembled *Aspergillus oryzae* genome FASTA files downloaded from NCBI using NCBI Datasets on 7 May 2026. Assembly quality was assessed using BUSCO with the aspergillus_odb12 lineage dataset before downstream annotation [16]. The 123 assemblies retained for the final analysis had complete BUSCO scores ranging from 98.7% to 99.5%. Three assemblies, Ao_TK-54, Ao_RIB326, and Ao_GCA_039654725, had lower BUSCO completeness scores of 95.5%, 95.9%, and 97.7%, respectively, and were excluded before ANNEVO annotation and EukPan processing. Individual complete BUSCO scores and post-screening inclusion status for all 126 assemblies are provided in Supplementary Table S1.

Genome assemblies were annotated using ANNEVO v2.2.3 [12], which predicts structural gene models directly from genome FASTA sequences without requiring matched RNA-seq data. The resulting GFF/GTF annotation files and their corresponding genome FASTA files were used as input to EukPan. EukPan v0.1.0 was run using genome files in the assembled_genomes directory, annotation files in the gff directory, pangenome_results as the output directory, and 32 threads.

Group labels followed the A-H classification defined by Watarai et al. [14]. The final dataset included 33, 5, 18, 2, 16, 13, 9, and 3 strains assigned to groups A-H, respectively; the remaining 24 strains were unassigned. Industrial-use metadata were taken from Watarai et al. [14]. The Japanese product names sake, miso, shoyu, and mirin refer here to Japanese rice wine, fermented soybean paste, soy sauce, and sweet rice seasoning, respectively. Shoyu is reported as soy sauce throughout this manuscript and Supplementary Table S1. Use in soy sauce production was recorded at the sample level for the TK strains identified in that study; use information was classified as not available when no corresponding information was available in the metadata examined. Sample identifiers, genome accession numbers, complete BUSCO scores, group assignments, industrial-use metadata, and post-screening inclusion status for all 126 screened assemblies are provided in Supplementary Table S1.

EukPan is a post-annotation workflow. In the ANNEVO–EukPan configuration used here, ANNEVO generated structural gene predictions from assembled genome FASTA files, and EukPan standardized and processed the predicted gene models for pangenome analysis. Thus, the combined ANNEVO–EukPan workflow can begin with genome FASTA files alone when structural annotations are unavailable, whereas EukPan itself requires corresponding genome FASTA and GFF/GTF annotation files. EukPan can also process existing GFF/GTF annotations generated using other structural annotation workflows. The Methods subsections whose headings begin with ‘EukPan’ describe operations performed by the core pipeline. ModelTest-NG, IQ-TREE 3, FigTree, and SplitsTree App were applied separately to EukPan outputs, whereas PCA used an optional EukPan helper script.

### EukPan annotation standardization and proteome extraction

Within the core EukPan pipeline, input GFF/GTF annotation files were converted and standardized using AGAT v1.7.0 [17]. For genes with multiple transcript isoforms, EukPan used AGAT to retain the longest isoform as the representative transcript. Protein sequences were then extracted from the corresponding genome FASTA and standardized annotation files using gffread v0.12.9 [18]. Protein FASTA headers were simplified using SeqKit v2.13.0 [19] to ensure consistent proteome identifiers for the subsequent OrthoFinder analysis.

### EukPan orthogroup inference and core-alignment construction

Within the core EukPan pipeline, orthogroup inference for the 123 extracted proteomes was performed using OrthoFinder v3.1.4 [20] in a dedicated conda environment. Orthogroups present in all 123 strains were classified as core orthogroups. EukPan identified 11,245 core orthogroups, of which 10,538 were single-copy orthologues suitable for core-alignment construction.

For each single-copy core orthologue, EukPan extracted one protein sequence per strain and confirmed that the resulting orthogroup FASTA file contained exactly one sequence from each of the 123 strains. Protein sequences were aligned using MAFFT v7.526 [21], and the resulting alignments were trimmed using trimAl v1.5.rev1 with the automated1 option [22]. No taxa were lost from any alignment during trimming. Three trimmed alignments shorter than 30 amino acids were excluded, leaving 10,535 trimmed single-copy core-orthologue alignments for concatenation. The final concatenated core-protein alignment was saved as concat_all.fa, together with an accompanying partition file.

### Downstream core-gene phylogeny and split-network analysis

The concatenated single-copy core-protein alignment generated by EukPan was used for downstream evolutionary analyses conducted outside the core EukPan pipeline. Amino acid substitution model selection was performed using ModelTest-NG v0.1.7 [23] with 32 threads (`modeltest-ng –i pangenome_results/phylo/concat_all.fa –d aa –p 32`).

To avoid repeating the computationally intensive model search in IQ-TREE, the selected VT+I+G4+F model was specified directly in IQ-TREE 3 v3.1.2 [24]. The concatenated alignment was analyzed under this single model without a partition scheme. A maximum-likelihood tree was inferred with automatic thread allocation, 1,000 ultrafast bootstrap replicates [52], and 1,000 SH-aLRT replicates [53] using the command `iqtree3 –s pangenome_results/phylo/concat_all.fa –st AA –m VT+I+G4+F –T AUTO –B 1000 –-alrt 1000 –-prefix`

`pangenome_results/phylo/iqtree_concat_VT+I+G4+F`.

The resulting maximum-likelihood phylogeny was visualized using FigTree v1.4.4 and midpoint-rooted for display in Fig. 2.

The same concatenated alignment was analyzed using SplitsTree App v6.4.16 [25] with uncorrected p-distances and the default NeighborNet procedure. In the saved SplitsTree project, the SPLITS block reported nsplits=156, and the network properties were recorded as cyclic with fit=98.4. These values were read directly from the saved NeighborNet output and were not estimated separately. For Supplementary Fig. S2, the maximum-likelihood tree was re-rooted on the branch leading to group A solely to facilitate visual comparison with the accessory-genome clustering. Group A was not treated as an evolutionary outgroup.

### EukPan shared accessory-genome processing and output-based analyses

Within the core EukPan pipeline, shared accessory orthogroups were identified from the OrthoFinder Orthogroups.GeneCount.tsv table. For orthogroup i and strain j, presence was defined as Iij = 1 when the corresponding gene count was greater than zero and Iij = 0 otherwise. The prevalence of orthogroup i was calculated as the row sum of Iij across the 123 strains.

Orthogroups present in all 123 strains were classified as core orthogroups. Non-core orthogroups present in at least two but fewer than all strains were classified as shared accessory orthogroups, corresponding to a prevalence range of 2–122 strains. Orthogroups detected in only one genome, here termed single-genome orthogroups, were excluded from the shared-accessory matrix and all subsequent accessory-genome analyses.

This minimum prevalence threshold was adopted because the study focused on recurrent gene-content variation among closely related strains rather than on compiling an exhaustive inventory of genes detected uniquely in individual genomes. In addition, unlike compact, intron-free prokaryotic coding sequences, eukaryotic gene calls depend on exon–intron prediction, transcript-model resolution, and representative-isoform selection. An orthogroup apparently restricted to a single eukaryotic genome may therefore reflect assembly fragmentation, missed or split gene models, differences in splice-model prediction, or isoform-processing effects rather than a genuine biological difference. Requiring detection in at least two independent genome assemblies provided a minimum level of recurrence before an orthogroup contributed to the comparative presence–absence matrix.

This EukPan filtering step yielded 4,407 shared accessory orthogroups. Only these orthogroups were used for the prevalence distribution, Jaccard-distance calculations, hierarchical clustering, heatmap construction, and PCA. Internal validation confirmed that all 4,407 rows in the shared-accessory matrix had prevalence values between 2 and 122 strains.

For the present study, downstream accessory-genome summary statistics were calculated directly from the validated binary matrix. For each orthogroup, prevalence was calculated as its row sum, and the frequency at prevalence k was defined as the number of orthogroups with a row sum of k. The numbers of orthogroups present in 10 or fewer strains and in 120 or more strains were calculated by summing the frequencies over k = 2–10 and k = 120–122, respectively. The counts reported at prevalence values of 2, 90, and 122 strains were read directly from this frequency distribution.

For each strain, the number of shared accessory orthogroups was calculated as the column sum of the binary matrix. The reported range corresponded to the minimum and maximum column sums, and the reported mean was the arithmetic mean across all strains.

Using the validated EukPan binary matrix, pairwise Jaccard distances between strains were calculated as one minus the ratio of the number of orthogroups present in both strains to the number present in either strain. Mean within-group distances were calculated using each unique unordered pair once, whereas the mean distance between group A and the non-A strains was calculated from all cross-group pairs. Hierarchical clustering was performed using average linkage with optimal leaf ordering. The resulting strain order was used to display the reordered Jaccard-distance matrix and the accessory-orthogroup presence–absence heatmap.

### Directional group-associated orthogroup extraction and functional characterization

Directional orthogroup sets were extracted from the EukPan gene-count output using the optional extract_group_presence_absence.py script. For each focal group, the non-group set comprised every other strain in the 123-genome analysis, including strains assigned to other A-H groups and the 24 unassigned strains. A group-associated orthogroup was defined as present in 90-100% of the focal group and 0-10% of the non-group set. A group-depleted orthogroup was defined using the reciprocal thresholds of 0-10% in the focal group and 90-100% in the non-group set. Presence was scored when the OrthoFinder gene count was greater than zero. Within these directional sets, strict denotes an all-versus-none distribution: 100% versus 0% for a strict group-associated orthogroup or 0% versus 100% for a strict group-depleted orthogroup. This is an operational prevalence label, not a separate orthology class. Because these contrasts were generated from the same shared-accessory analysis, single-genome orthogroups remained excluded.

For each directional set, the optional extract_representative_sequences.py script selected the longest protein sequence among all members of each orthogroup. Functional annotation was performed using OmicsBox v4.0.59 (build 5da612bb05; BioBam Bioinformatics, Valencia, Spain), and protein domains and functional sites were predicted using InterProScan v5.78-109.0 [54] against InterPro release 109.0. Sequence-similarity searches used DIAMOND blastp v2.2.2.182 [55] against the NCBI non-redundant (nr) protein database released on 2 July 2026, with standard sensitivity and an E-value threshold of 1.0 x 10^-15^. Searches were taxonomically restricted to *Aspergillus oryzae* (Taxonomy ID 5062), *Aspergillus sojae* (Taxonomy ID 41058), and *Aspergillus tamarii* (Taxonomy ID 41984), except for the group-H-associated set (H_com), which was searched without a taxonomic filter because multiple sequences yielded no hit when the filter was applied.

Gene Ontology terms were mapped and assigned using the Blast2GO-based GO Annotation feature in OmicsBox [56] with default parameters: an E-Value-Hit-Filter of 1.0 x 10^-6^, an annotation cutoff of 55, and default evidence-code weights. Orthogroup identifiers in the presence-absence tables, representative-protein FASTA files, and OmicsBox exports were cross-checked and matched across all 16 directional sets.

For functional characterization, GO identifiers in the OmicsBox GO and InterPro-derived GO fields were combined and deduplicated within each orthogroup; InterPro identifiers were likewise deduplicated within each orthogroup. Annotation coverage was calculated as the proportion of orthogroups with at least one GO term or InterPro identifier. Term frequency was calculated as the number of orthogroups carrying a term divided by the total number of orthogroups in the corresponding directional set. These frequencies were used to describe the functional composition of the lists; no statistical hypothesis testing was performed and no P values were calculated. The complete directional orthogroup lists, prevalence values, representative-sequence identifiers, descriptions, GO terms, InterPro identifiers, and enzyme annotations are compiled in Supplementary Table S2.

### Evidence-filtered protein homology and literature review

Representative proteins from the group-A-associated, group-A-depleted, group-E-associated, and group-E-depleted sets were searched against 5,612 reviewed *Aspergillus* proteins obtained from UniProtKB/Swiss-Prot on 29 August 2026 using blastp in NCBI BLAST+ v2.17.0 [57]. Searches used an E-value threshold of 1 x 10-5, –max_target_seqs 5, and SEG filtering. Reciprocal searches between associated and depleted representative proteins were also inspected for possible gene-model fragmentation or orthogroup splitting.

Evidence categories were assigned through a qualitative review of sequence-search results and the primary experimental literature, outside the automated EukPan pipeline, by integrating amino acid identity, query coverage, the specificity of the leading reviewed hits, and whether the corresponding homolog had been investigated by gene disruption or complementation, heterologous expression, biochemical assay, or coupled genetic-metabolite analysis. The high-evidence category required a near-full-length, high-identity match to a homolog with experimental support for its function. Up to five returned hits were examined, and the best sequence hit was distinguished from the homolog supporting the experimental interpretation when these differed. For OG0012552, this distinction separates the best AflL hit from the experimentally investigated StcL homolog; the supporting alignments and literature links are provided in Supplementary Data S1. The category was not an automatic EukPan, OmicsBox, or InterProScan output, nor a statistical score. Only high-evidence candidates were linked to experimentally characterized homologs in the Results and Discussion; lower-confidence OmicsBox and InterPro annotations were described only as hypotheses and were not treated as proof of biochemical activity.

### Genomic comparison of strict group-A-depleted orthogroups between group-F RIB40 and group-A TK-34

RIB40, assigned to group F (GenBank assembly GCA_000184455.3), served as the coordinate and annotation reference, whereas TK-34, assigned to group A (GCA_009686785.1), served as the representative group-A comparison assembly. The strict group-A-depleted subset comprised orthogroups absent from all 33 group-A strains and present in all 90 non-A strains. Representative proteins were mapped by BLASTP to the ANNEVO-derived RIB40 proteome. A mapping was classified as high confidence when E <= 1 x 10^-20^, query coverage was at least 80%, and identity was at least 70%; a mapping was retained as usable when E <= 1 x 10^-10^, query coverage was at least 50%, and identity was at least 30%. All 93 candidates mapped to RIB40 (87 high-confidence and six usable mappings). Candidate genes were joined into a compact cluster when successive genes were separated by no more than two intervening genes and no more than 20 kb. Gene labels used established common names when supported by high-category evidence; otherwise the gene, product, or locus_tag field from the RIB40 RefSeq GenBank record GCF_000184455.2 was used.

For nucleotide-level comparison, each RIB40 candidate cluster plus 20 kb on each side was searched against the TK-34 genome with blastn in megablast mode. Searches used E <= 1 x 10^-20^, minimum nucleotide identity 90%, dust filtering disabled, max_target_seqs 100,000, and max_hsps 10,000. Alignment blocks of at least 500 bp were used to evaluate local collinearity. An operational two-flank deletion relative to RIB40 was called when the candidate interval was overlapped by an unaligned RIB40 segment bracketed by collinear alignments on the same TK-34 contig, the TK-34 interflank span was no more than 500 bp, the inferred missing RIB40 segment was at least 1 kb, and at least 80% of the candidate-cluster span overlapped the unaligned interval. Small negative interflank spans denote overlapping BLAST alignments. These comparisons identify RIB40 sequence absent at the corresponding TK-34 locus; they do not polarize the evolutionary event as a loss in TK-34 rather than a gain in RIB40. Regions lacking both flanks on one TK-34 contig were classified as unresolved, and regions with substantial sequence between the TK-34 flanks were classified as partial/complex or replacement-like.

Chromosome-end proximity was evaluated from the RIB40 genome FASTA sequence lengths and the mapped cluster coordinates. For a cluster spanning coordinates s through e on a chromosome of length L, distances from the left and right chromosome ends were calculated as s – 1 and L – e, respectively. C6 spanned positions 5,051,763-5,065,945 on chromosome 3 (L = 5,123,684 bp), placing its boundaries 57,739-71,921 bp from the right end. C7 spanned positions 4,859,363-4,866,494 on chromosome 4 (L = 4,887,096 bp), placing its boundaries 20,602-27,733 bp from the right end. C12 spanned positions 25,245-61,362 on chromosome 7 (L = 2,933,481 bp), placing the complete interval within 61,362 bp of the left chromosome end. Because each complete interval was within 100 kb of a chromosome end, C6, C7, and C12 were classified as subtelomeric following the operational interval used in the comparative section *Flavi* analysis [35].

### PCA using an optional EukPan helper script

PCA was performed using an optional EukPan helper script and the shared accessory-orthogroup presence–absence matrix. The matrix was transposed so that rows represented strains and columns represented orthogroups. Orthogroup columns were centered but were not standardized to unit variance, and zero-variance columns were removed before analysis. Principal component scores were calculated by singular value decomposition.

For component k, the percentage of explained variance was calculated as 100 multiplied by the squared singular value of component k and divided by the sum of the squared singular values across all components. Score ranges were defined as the observed minimum and maximum values within each labeled set. The separation gap along PC1 was calculated as the minimum PC1 score among group-A strains minus the maximum PC1 score among all non-A strains. Group labels were assigned after PCA for visualization and comparison and were not used to calculate the principal components.

### Use of generative AI tools

ChatGPT (OpenAI) was used for English-language revision of the manuscript and for programming assistance during the development and validation of analysis scripts. The authors reviewed and edited all AI-assisted outputs and take full responsibility for the final text, code, analyses, interpretations, and conclusions.

## Data availability

The genome assemblies screened in this study are publicly available from NCBI; their accession numbers are listed in Supplementary Table S1. The EukPan shared accessory-orthogroup tables and compact downstream outputs supporting Figs. 3-6 are provided as Supplementary Data S1. Directional group-associated and group-depleted orthogroup lists, together with their OmicsBox GO and InterPro annotations, are provided in Supplementary Table S2. The dedicated ‘Strict A-depleted’ worksheet in Supplementary Table S2 lists all 93 strict group-A-depleted orthogroups with group and non-group prevalence, RIB40 coordinates and annotations, compact-cluster assignments, and TK-34 comparison classifications. The underlying high-evidence protein-homology summary and supporting BLASTP hits, nucleotide-alignment table, cluster-level summary, and machine-readable strict-orthogroup table supporting Fig. 8 are included in Supplementary Data S1, together with all 16 representative-protein FASTA files, the core-protein alignment, and the original IQ-TREE 3 tree. The alignment is supplied in losslessly reformatted interleaved NEXUS format with per-sequence checksums.

## Code availability

The EukPan source code, documentation, and included downstream helper scripts are available at https://github.com/yukio-nagano/eukpan and are archived as EukPan v0.1.0 in Zenodo (https://doi.org/10.5281/zenodo.22093294; commit 40b5dd35075b1b36ac1ac16cd4b7cb9170a45aae).

## Acknowledgements

We thank Hisanori Tamaki of the Faculty of Agriculture, Kagoshima University, and Genta Kobayashi of the Faculty of Agriculture, Saga University, for their support and collaboration in this IFO-supported study.

## Funding

This study was supported in part by the Institute for Fermentation, Osaka, Japan (IFO; grant number LA-2024-008).

## Author contributions

K.S. and Y.N. conceived and designed the study, performed the bioinformatic analyses, and interpreted the data. M.G. and T.F. performed detailed analyses of the *Aspergillus* comparative genomic results. Y.N. wrote the original manuscript draft, and all other authors reviewed and edited the manuscript. All authors read and approved the final manuscript.

## Competing interests

For transparency, Yukio Nagano conducts collaborative research with Senoo Suisan, Inc., and Sante Laboratories, Inc., and serves as a director of Smart Review Technologies Co., Ltd. These relationships are independent of the present study. All authors declare no competing interests in relation to this study.

## Notes

### Competing Interest Statement

The authors have declared no competing interest.

