## Supplementary Information for "The automated eukaryotic pangenome pipeline EukPan reveals accessory genome differentiation beyond core-gene phylogeny in *Aspergillus oryzae*"

#### Contents

**Supplementary Figure S1**. NeighborNet split network based on core-protein sequences

**Supplementary Figure S2**. Alternative display of the core-gene phylogeny

**Supplementary Figure S3**. Numbers of group-associated and group-depleted orthogroups for groups A-H

**Supplementary Figure S4**. Functional-annotation coverage across the 16 directional sets

### Supplementary Figure S1

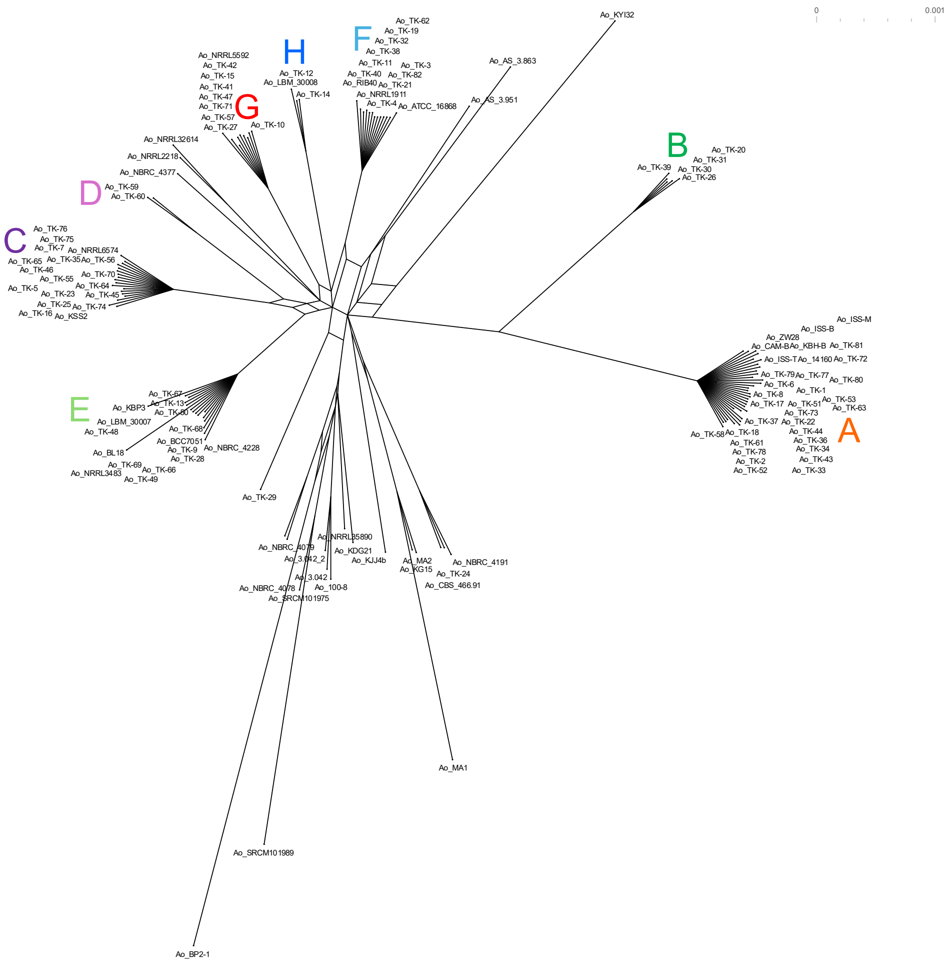


**Supplementary Figure S1.** NeighborNet split network based on core-protein sequences. The network was constructed using SplitsTree App from uncorrected p-distances calculated from the same concatenated core-protein alignment used for Fig. 2. The saved NeighborNet output contained 156 splits, was classified as cyclic, and had a reported fit value of 98.4%. Box-like structures indicate conflicting phylogenetic signals within the alignment. Colored labels indicate the previously defined groups A-H.

### Supplementary Figure S2

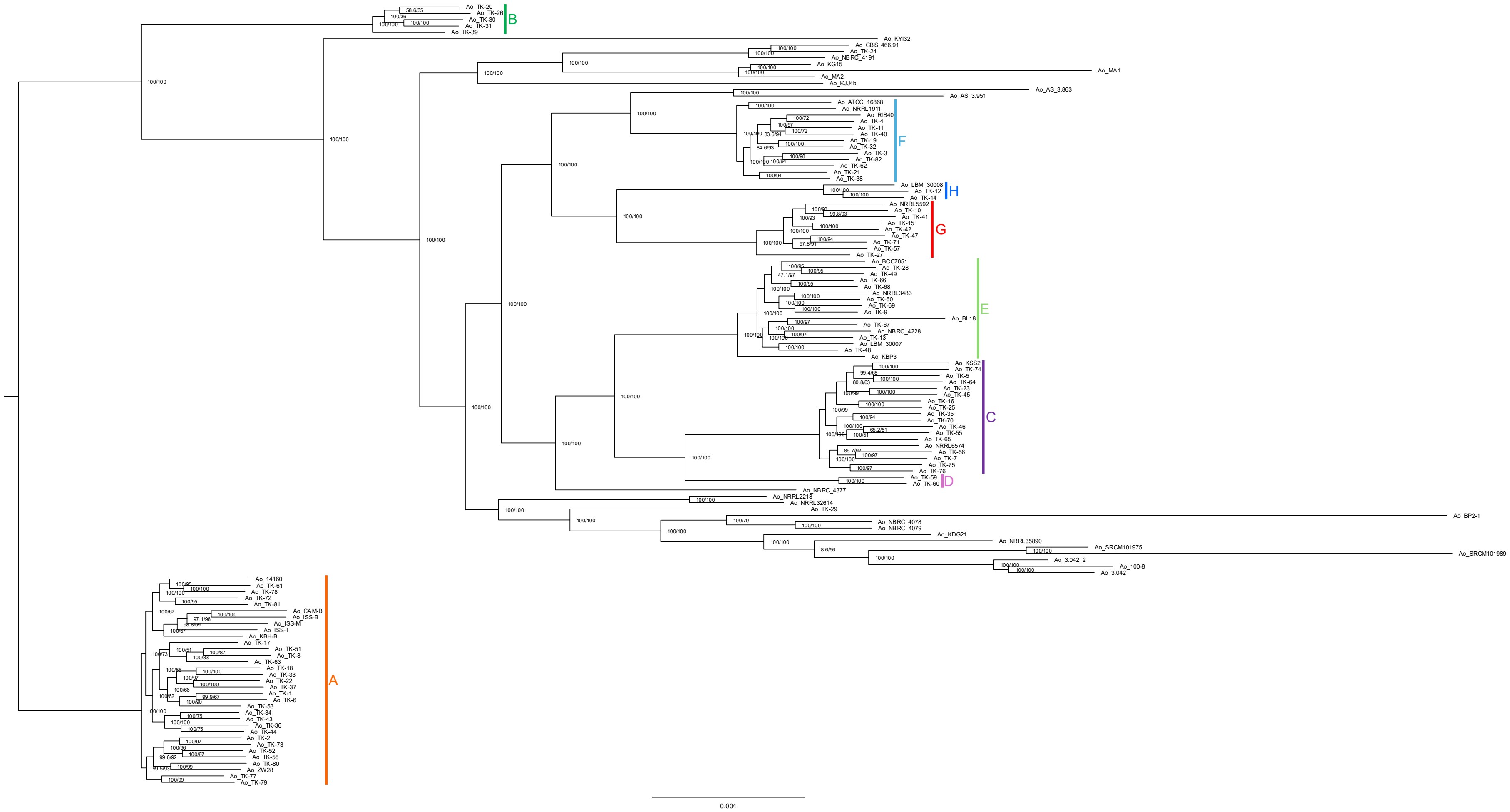


**Supplementary Figure S2.** Alternative display of the core-gene phylogeny. The maximum-likelihood tree shown in Fig. 2 was re-rooted on the branch leading to group A solely to facilitate visual comparison with the accessory-genome clustering. Group A was not treated as an evolutionary outgroup. Colored brackets indicate the previously defined groups A-H.

#### Supplementary Figure S3

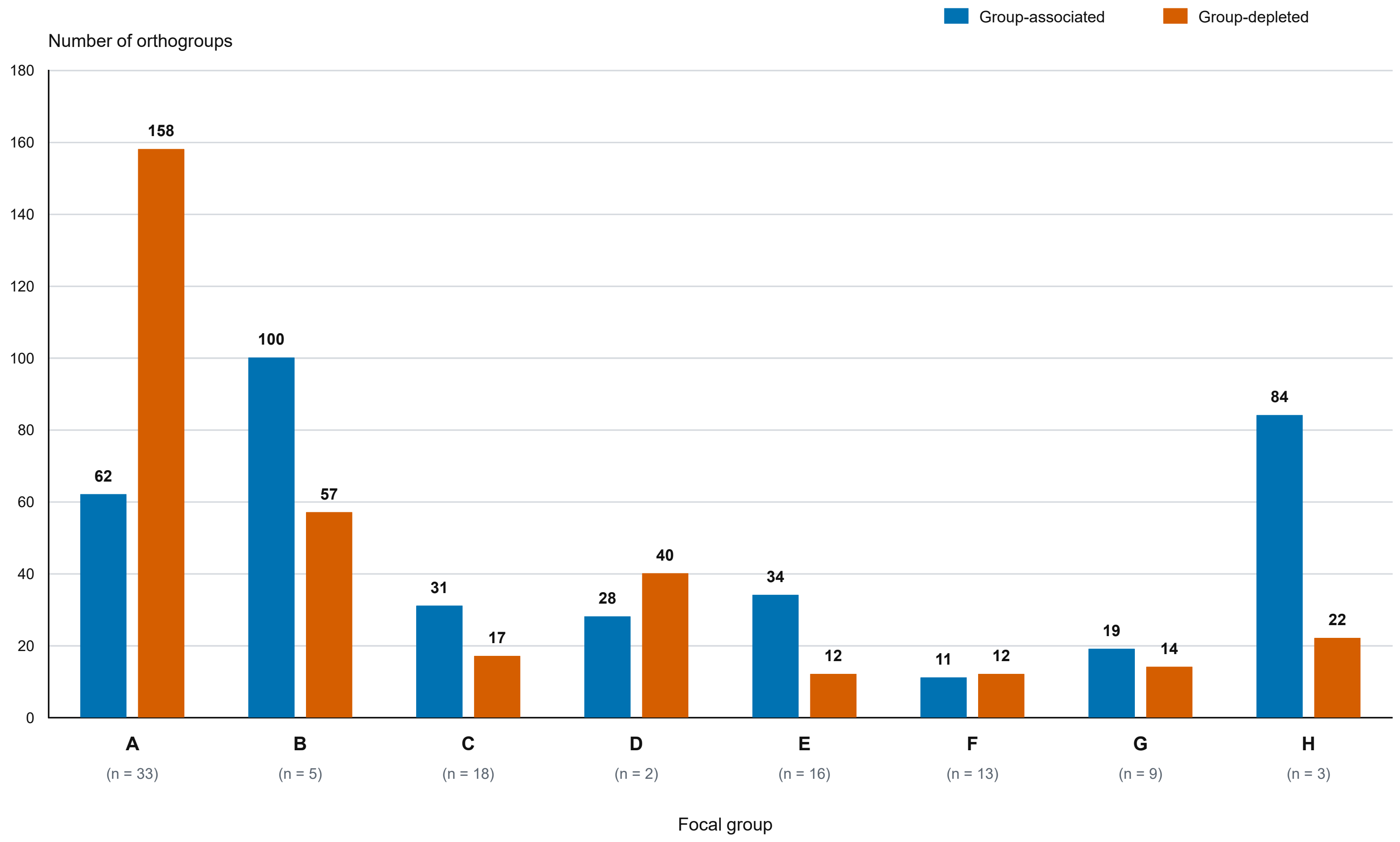


**Supplementary Figure S3.** Numbers of group-associated and group-depleted orthogroups for groups A-H. Bars show orthogroup counts for each directional set, and focal-group sample sizes are shown below the group labels. Group-associated orthogroups were present in 90-100% of the focal group and 0-10% of the complementary set; group-depleted orthogroups met the reciprocal thresholds.

### Supplementary Figure S4

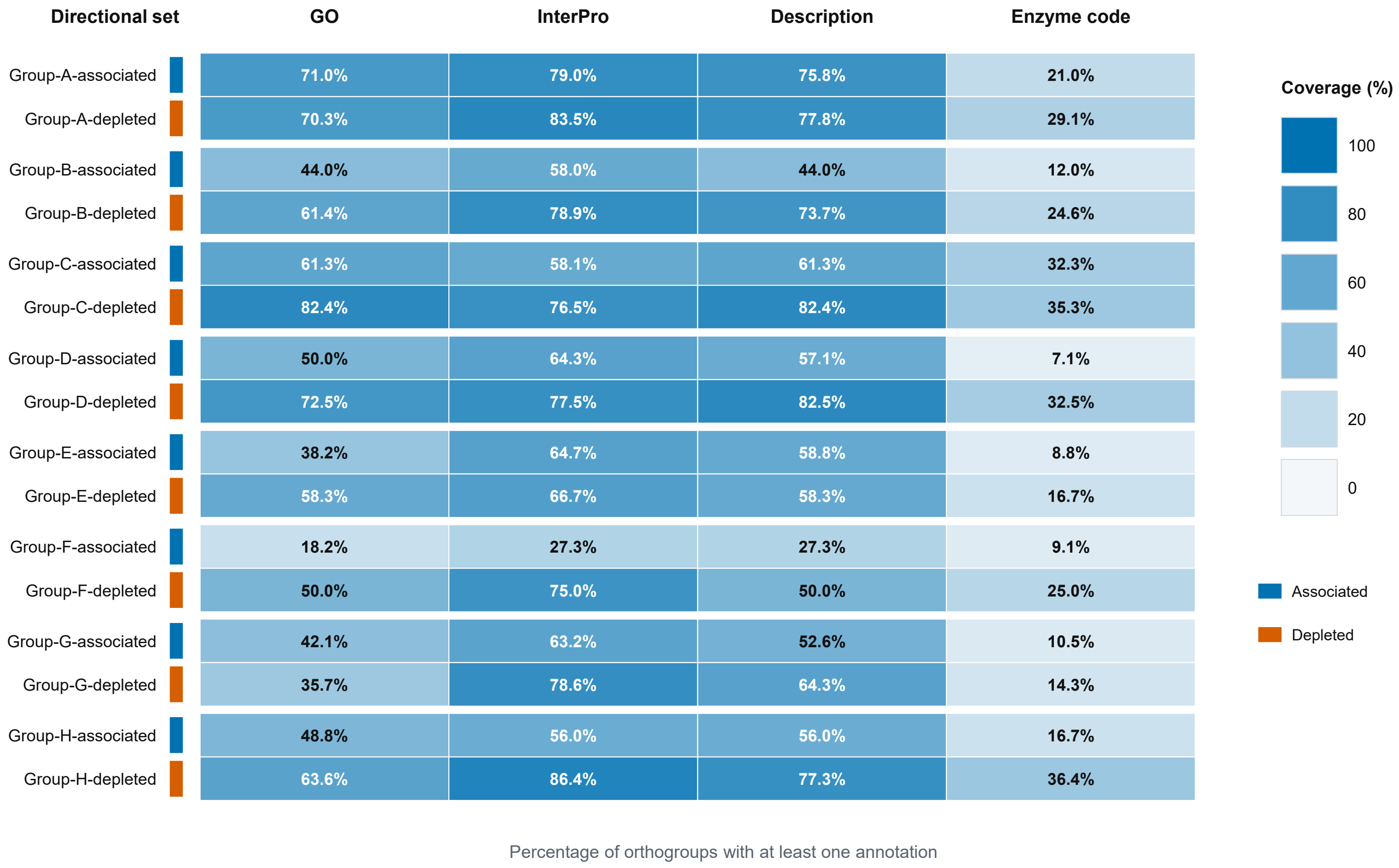


**Supplementary Figure S4.** Functional-annotation coverage across the 16 directional sets. Cell values show the percentage of orthogroups with at least one GO term, InterPro identifier, informative sequence-similarity description, or enzyme code. GO coverage uses the union of OmicsBox and InterPro-derived GO identifiers. Blue and orange row markers indicate group-associated and group-depleted sets, respectively. These values describe annotation coverage; no statistical enrichment testing was performed.
