## Supplementary Data S1 for "The automated eukaryotic pangenome pipeline EukPan reveals accessory genome differentiation beyond core-gene phylogeny in *Aspergillus oryzae*": README.txt

Supplementary Data S1
EukPan core- and accessory-genome source data for the 123-genome Aspergillus oryzae analysis

This archive contains tabular EukPan outputs and compact downstream tables used
to calculate or reproduce the accessory-genome results reported in Figs. 3-6
and the RIB40-TK-34 locus comparisons reported in Fig. 8. Shared accessory
orthogroups are non-core orthogroups detected in 2-122 of the 123 analyzed
samples. Orthogroups detected in only one sample are outside the scope of these
files.

For Fig. 8, a strict group-A-depleted orthogroup was operationally defined as
detected in 0/33 group-A strains and 90/90 non-A strains. The term strict denotes
this all-versus-none prevalence criterion and not a separate orthology class.
RIB40 is assigned to group F and TK-34 to group A; genome accessions and group
assignments are provided in Supplementary Table S1.

Contents

accessory_results/
  shared_accessory_gene_count.tsv
    Gene-count table for the 4,407 shared accessory orthogroups.
  shared_accessory_presence_absence.tsv
    Binary presence/absence table used for accessory-genome analyses.
  shared_accessory_summary.tsv
    Orthogroup-level summary table.

accessory_analysis/
  00_summary.tsv
    Dataset and per-sample accessory-orthogroup counts.
  01_presence_distribution.tsv
    Number of orthogroups at each prevalence across the 123 samples.
  03_jaccard_distance_matrix.tsv
  03_jaccard_distance_matrix_reordered.tsv
  03_jaccard_similarity_matrix.tsv
  03_jaccard_similarity_matrix_reordered.tsv
    Pairwise Jaccard distance and similarity matrices used for Figs. 5 and 6.

accessory_pca/
  accessory_presence_absence_matrix_for_R.tsv
    Presence/absence matrix supplied to the PCA workflow.
  accessory_pca_metadata.tsv
    Sample labels and group metadata used for visualization.
  accessory_pca_scores.tsv
    PCA scores for the analyzed samples.
  accessory_pca_used_orthogroups.tsv
    Orthogroups retained in the PCA input matrix.
  accessory_pca_variance.tsv
    Explained-variance results for the principal components.

Fig_8_source_data/
  strict_group_A_depleted_orthogroups.tsv
    Machine-readable list of all 93 strict group-A-depleted orthogroups, with
    RIB40 coordinates, annotations, compact-cluster assignments, and TK-34
    comparison classifications. The same records are presented in the
    'Strict A-depleted' worksheet of Supplementary Table S2.
  high_evidence_functional_matches.tsv
    Eight high-evidence reviewed-protein matches used for functional discussion.
  A_depleted_RIB40_TK34_deletion_summary.tsv
    Coordinates and TK-34 comparison classifications for the 15 RIB40 clusters.
  A_depleted_RIB40_official_annotations.tsv
    RIB40 mapping and RefSeq annotation records for all 93 strict candidates.
  RIB40_regions_vs_TK34_megablast.tsv
    Nucleotide alignments used to evaluate local collinearity and deletion calls.

pipeline.log
  EukPan run log recording input counts, processing steps, software checks, and
  final output counts.

Large raster/vector figure files, EMF files, and the full PCA loading table were
not duplicated in this archive to limit file size. Public genome accessions are
listed in Supplementary Table S1. The included binary matrices, source tables,
and alignment records are sufficient to reconstruct the reported prevalence,
Jaccard-distance, hierarchical-clustering, heatmap, PCA, and Fig. 8 summaries.

EukPan source code and documentation
https://github.com/yukio-nagano/eukpan

phylo/
  iqtree_concat_VT+I+G4+F.treefile
    Original IQ-TREE 3 Newick output, including support labels and branch lengths.
  concat_all_interleaved.nex
    The exact concatenated core-protein alignment (123 samples, 5,009,064 sites),
    reformatted from concat_all.fa as interleaved NEXUS in blocks of 1,000 sites
    to improve ZIP compression. Sample order, sequences, gaps, and site order
    are unchanged. The original analysis used concat_all.fa.
  aligned_sequence_checksums.tsv
    SHA-256 of each full aligned amino acid string, excluding line breaks,
    for verifying lossless conversion.

directional_representative_proteins/
  Sixteen original representative-protein FASTA files used for the directional
  group analyses and OmicsBox annotation. Filenames containing common_non identify
  group-associated sets; rare_non identifies group-depleted sets.

GO coverage and term frequencies use the union of OmicsBox GO and InterPro-derived
GO identifiers, deduplicated within each orthogroup. The original two annotation
fields are retained separately in Supplementary Table S2.

phylo/core_network_blocks.nex
  Saved TAXA, DISTANCES and SPLITS blocks from the SplitsTree project.
  The redundant input TAXA block and workflow LINK directives were omitted.
  Split weights, distances, fit=98.4 and nsplits=156 are unchanged.

Reference verification (13 September 2026): high_evidence_functional_matches.tsv distinguishes the best reviewed hit from the literature-supported homolog for AflL/StcL, OmtB/DmtA, and OmtA. The added high_evidence_candidates_original_blastp.tsv contains unmodified original hits for the eight candidates. Columns: qseqid, sseqid, pident, length, mismatch, gapopen, qstart, qend, sstart, send, evalue, bitscore, qlen, slen, qcovs, stitle. No BLAST searches or pangenome analyses were rerun. High denotes homology-based support, not direct functional validation in the analyzed strain.
